# Characterization of active kinase signaling pathways in astrocytes and microglia

**DOI:** 10.1101/2025.04.18.649617

**Authors:** William G. Ryan, Hunter M. Eby, Nicole R. Bearss, Ali S. Imami, Abdul-rizaq Hamoud, Priyanka Pulvender, Justin L. Bollinger, Eric S. Wohleb, Robert E. McCullumsmith

**Affiliations:** Department of Neurosciences and Psychiatry, College of Medicine and Life Sciences, University of Toledo, Toledo, OH, USA; Department of Pharmacology & Systems Physiology, College of Medicine, University of Cincinnati, Cincinnati, OH, US; Neurosciences Institute, ProMedica, Toledo, OH, USA

**Keywords:** *kinase network*, *active kinome*, *active signaling network*, *astrocyte*, *microglia*, *systems biology*

## Abstract

Protein kinases are central to healthy brain function, regulating critical cellular processes through complex signaling networks. However, understanding differences in kinase signaling of brain cells remains a preeminent challenge of neuroscience. This study aimed to characterize kinase pathways enriched in astrocytes and microglia isolated from male and female murine prefrontal cortex. Using the PamGene PamStation®12 platform, we discovered cell-type-specific kinomic profiles and computationally reconstructed each cell type’s unique active signaling protein-protein interaction network. Notably, our analysis revealed minimal overlap between kinase activity and respective cell-subtype specific kinase transcriptional profiles identified in the Allen Mouse Whole Brain Transcriptomic Cell Type Atlas, highlighting an important limitation of relying solely on gene mRNA expression levels for functional inference in kinase focused studies. These findings also suggest that cell- and sex-specific protein kinase signaling may influence susceptibility to deleterious brain conditions and consequently underscore the importance of considering activity as a biological variable in systems research, offering a new framework for developing targeted therapeutic interventions in precision medicine.

## Introduction

Protein kinases are critical regulators of a multitude of cellular processes, acting as influential nodes in signaling networks that control cellular growth, differentiation and survival [1]. Protein kinases function by transferring phosphate groups from ATP to specific substrates, thereby modulating the activity, localization and interaction of target proteins [2]. In the brain, kinases are instrumental in orchestrating complex signaling pathways that underlie neuronal communication, synaptic plasticity and synaptic pruning responses to environmental stimuli [3–5]. Given these fundamental roles, it is not surprising that dysregulation of kinase activity is implicated in a variety of brain disorders, including Alzheimer’s disease, schizophrenia and major depression [6–10]. Consequently, kinases have emerged as promising targets for the development of therapeutic interventions aimed at restoring normal brain function [11]. Despite the recognized importance of kinases in brain health, understanding their precise roles has been challenging [12]. Traditionally, studies have relied on measurements of gene expression or total protein abundance to infer the activity of kinases and other signaling proteins. However, recent advances in systems biology have revealed significant limitations in these approaches [13–15]. Notably, it has become clear that mRNA levels often do not correlate well with protein abundance [16, 17]. However, even when proteins are abundantly expressed, their activity may not be directly inferred from their expression levels [18]. This disconnect arises because protein function is not solely determined by its abundance, but also post-translational modifications, protein-protein interactions with other molecules, as well as subcellular localization [19].

Given these challenges, there has been a growing shift toward directly assessing kinase activity to gain a more accurate understanding of cellular signaling networks [20]. Active kinome profiling, which focuses on identifying kinases that are functionally active and involved in signaling networks, has become an increasingly valuable approach [21]. This method allows researchers to capture the dynamic state of signaling networks, offering insights not accessible through transcriptomic or proteomic analyses [22]. By identifying active kinases and their associated active signaling pathways, researchers can uncover the molecular mechanisms that drive cellular behavior in both normal and diseased states [23].

In the brain, astrocytes and microglia play pivotal roles in maintaining neuronal function and responding to injury or disease [24]. Astrocytes are involved in neurotransmitter recycling, blood-brain barrier maintenance and modulation of synaptic activity [25], while microglia act as the resident immune cells of the brain, mediating inflammatory responses and clearing cellular debris [26]. Despite their importance, the active signaling networks within these cell types remain poorly understood, particularly in the context of differences that may influence brain function and disease susceptibility [27–29]. To our knowledge, this study is the first to apply the PamGene PamStation®12 kinome activity profiling platform [30] to profile astrocytes and microglia isolated from murine prefrontal cortex. The kinome array allows for the simultaneous measurement of kinase activity across a wide array of substrates, providing a comprehensive view of the active kinome [31]. We characterized cell- and sex-specific differences in kinase activity and active signaling pathways of astrocytes and microglia, which has important implications for understanding the molecular basis of deleterious brain conditions as well as developing targeted therapies for these difficult to treat disorders.

## Materials and Methods

### Enrichment of astrocytes or microglia with fluorescence activated cell sorting (FACS)

Work was done in accordance with National Institute of Health guidelines for Care and Use of Animals and the University of Cincinnati Institutional Animal Care and Use Committee. Female C57BL/6J (n=3, Jackson #000664) and male DBA (n=3, Jackson #000671) mice, aged 6.5 to 7.5 weeks, were housed under standard conditions with ad libitum access to food and water. Following euthanasia by cervical dislocation, prefrontal cortex (PFC) tissue was dissected, and cells processed for astrocyte and microglia isolation. Astrocytes were enriched using a Percoll gradient, followed by staining with FITC-CD11b and PE-Recombinant-ACSA2 antibodies, as previously described [32]. Microglia were similarly enriched using a Percoll gradient and stained with PerCP-Cy5.5 Rat Anti-CD11b and PE-CF594 Rat Anti-Mouse CD45 antibodies, as previously described [33]. Cell suspensions were analyzed and sorted using a BioRad S3e four-color cytometer.

### Identification of differentially phosphorylated peptides on the PamGene kinome array

Cell pellets from astrocytes and microglia cultures were lysed using M-PER buffer containing Halt Protease and Phosphatase Inhibitor Cocktail. Lysates were centrifuged, and supernatants were assayed for protein concentration using the Pierce BCA Protein Assay Kit. Samples were diluted to 1 µg/µL and stored at −80°C, with frozen aliquots used only once to prevent loss of kinase activity. Kinase activity profiling was performed using the PamGene PamStation12 microarray as previously described [34], with 2 µg of protein loaded per well onto the STK PamChip®4. Phosphorylation was monitored in real-time using FITC-labeled anti-phospho peptide antibodies, and the peptide phosphorylation intensity was captured and analyzed using PamGene provided BioNavigator software. Peptides with differential changes in phosphorylation (≥ 15% change) were identified as previously described using the KRSA (Kinome Random Sampling Analyzer) software [35].

### Inference of upstream active protein kinases

Kinase Enrichment Analysis 3 (KEA3) [36] was used to identify upstream active kinases responsible for the observed differentially phosphorylated peptides. Mean p-values and kinome rank scores were computed using one-sided Fisher’s Exact Tests across KEA3’s kinase-substrate interaction libraries, including PhosphoSitePlus, PTMsigDB, and the ‘Cheng et al.’ library, which aggregates data from Phospho.ELM, HPRD, PhosphoNetworks, and PhosphoSitePlus. 1,000 bootstrap iterations were performed by randomly sampling the same number of peptides identified in each group comparison. This bootstrapping generated an expected mean rank and rank variance for each kinase, from which Enrichr combined scores were calculated as previously described [37]. The top 10% of kinases based on this combined score were selected as active kinases as previously described [38].

### Determination of cell-type specific kinase gene expression in mouse whole cortex and hippocampus

Allen Brain Map Mouse Whole Cortex and Hippocampus 10x single-cell transcriptome reference atlas [47] data was obtained as normalized trimmed means of gene expression aggregated per cell type. The kinome as defined previously by Moret *et al.* [48] was used to identify kinases expressed by astrocytes or microglia compared to other cell types for overlap with active kinases.

### Inference of upstream enriched transcription factors

ChIP-X Enrichment Analysis 3 (ChEA3)’s [39] brain-specific transcription factor (TF) library was used to infer TFs that are upstream of the differentially phosphorylated peptides and active kinases in each group. P-values and ranks for each TF were calculated using one-sided Fisher’s Exact Tests with CHEA3’s brain-specific transcription factor library. 1,000 bootstrap iterations were performed, and Enrichr combined scores were calculated to select the top 10% of TFs by combined score as enriched upstream TFs.

### Reconstruction of active signaling pathways with network-based integration

The Kinograte R software [40], which implements an optimized version of the well-established PCSF [41] algorithm, was used to generate active signaling protein-protein interaction (PPI) networks integrating previously identified peptides, kinases, transcription factors and algorithmically identified Steiner hidden nodes in each group as previously described [42]. Node prizes were assigned by percentile rank of peptide log2FoldChange or kinase and transcription factor combined scores returned from KEA3 and CHEA3 analyses. Edge costs were assigned by percentile rank of inverse STRING-DB [43] interaction confidence (minimum confidence .5) or phuEGO [44] semantic similarity. Over-representation analysis was performed using each PPI network’s nodes as input to the Enrichr [37] web app with the Gene Ontology database [45]. Enriched terms (FDR adjusted p-value < 0.05) were functionally clustered and visualized using PAVER (Pathway Analysis Visualization with Embedding Representations) [46], a meta-clustering method for pathways which identifies most representative terms (MRTs) for hierarchically clustered pathway embeddings by selecting whichever term is most cosine similar to its respective cluster’s average embedding. PAVER generated dot plots of mean cluster enrichment, cluster size and dissimilarity of the cluster MRT to its respective cluster’s average pathway embedding; Uniform Manifold Approximation Projection (UMAP) scatter plots of individual pathways colored by the cluster they belong to; and heatmaps showing enrichment of individual pathways in their identified cluster. Finally, HUGO Gene Nomenclature Committee (HGNC) symbols annotated to enriched pathways were used to generate informed subnetworks in order to visualize PPIs.

## Results

### Cell and sex specific kinase reporter phosphopeptide phosphorylation profiles

We first measured kinase activity in isolated astrocytes and microglia from the PFC of male and female mice (Figure 1). Using the PamGene kinome array, we identified unique phosphorylation profiles of serine/threonine reporter peptides (Figure 2A, 2B). In male microglia, 47 peptides were differentially phosphorylated, whereas in female microglia, 37 peptides were differentially phosphorylated; in male astrocytes, 48 peptides were differentially phosphorylated, whereas in female astrocytes, 38 peptides were differentially phosphorylated (Table S1). These findings indicated the possibility of distinct kinase activity patterns between astrocytes and microglia, as well as sex-specific differences in kinase activity.

**Figure 1.**
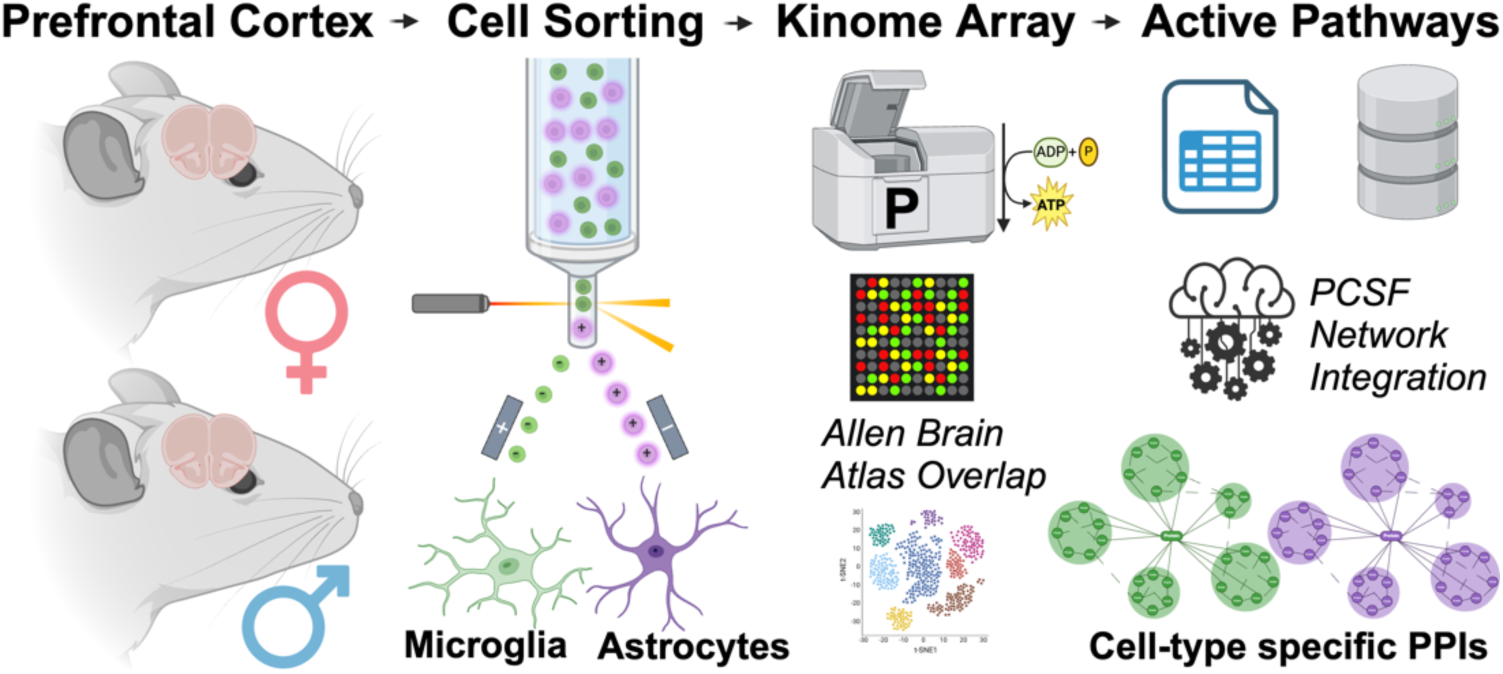
Overview of study design. Prefrontal cortex was harvested from male and female wildtype mice (n=3). Collected cells were enriched into populations of astrocytes or microglia with FACS. Differential kinase activity was profiled in each group versus their respective controls using the PamGene PamStation®12 PamChip®4. Active signaling pathways were reconstructed with PCSF network-based integration of differentially phosphorylated peptides, active kinases and enriched transcription factors. FACS: Fluorescence-activated cell sorting; PCSF: Prize-Collecting Steiner Forest.

**Figure 2.**
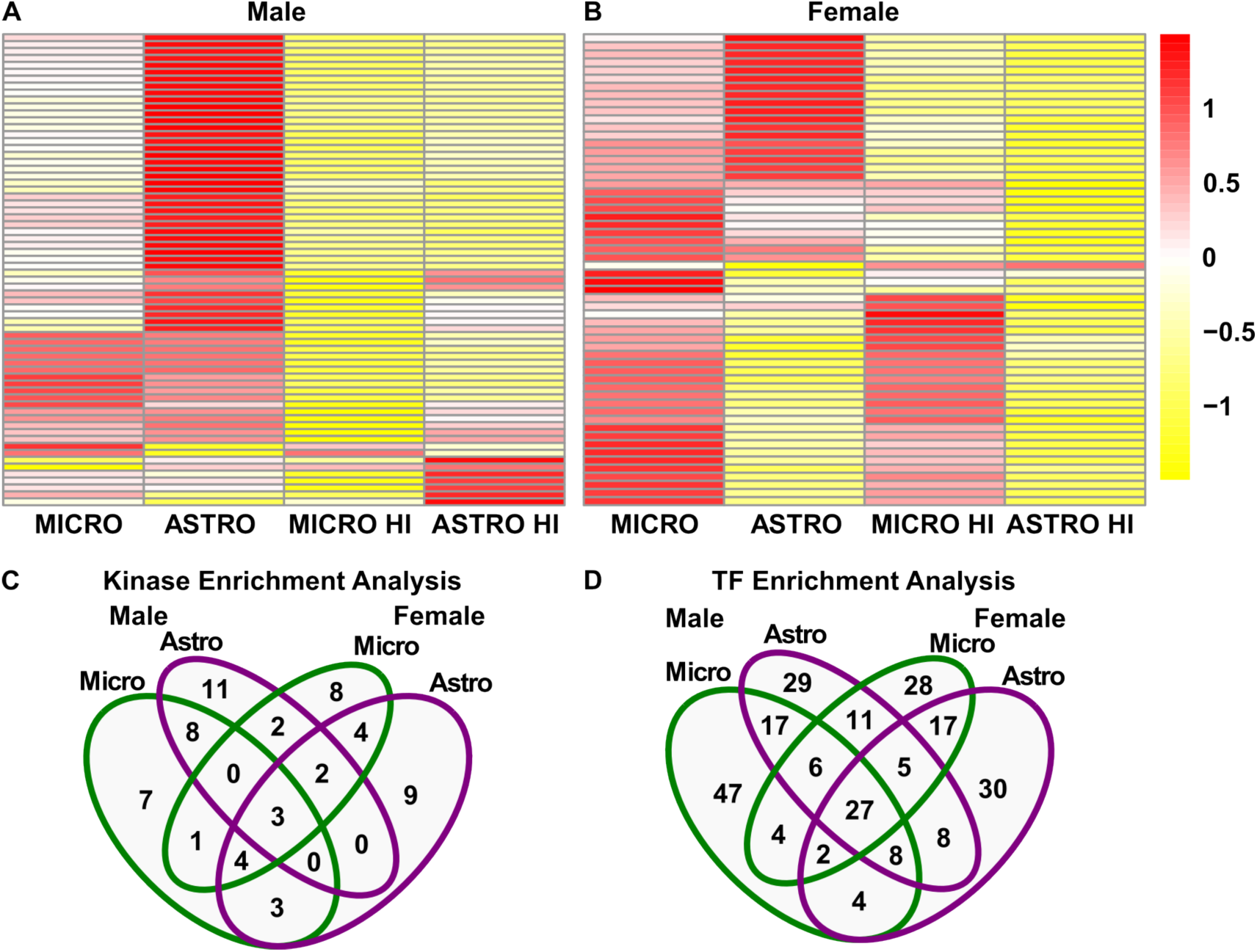
Prediction of upstream active kinases and enriched transcription factors associated with differentially phosphorylated peptides in microglia or astrocytes. Microglia or astrocytes from male and female wildtype mice PFC (n=3) were assayed on the PamGene PamStation®12 PamChip®4. Differentially phosphorylated reporter phosphopeptides were submitted to KEA3 to predict active kinases. Peptides and active kinases were submitted to CHEA3 to predict enriched transcription factors. **A.** Heatmap of mean normalized signal of differentially phosphorylated reporter phosphopeptides in male astrocytes and male microglia. **B.** Heatmap of mean normalized signal of differentially phosphorylated reporter phosphopeptides in female astrocytes and female microglia. **C.** Venn diagram overlap of identified active kinases. **D.** Venn diagram overlap of identified enriched transcription factors. HI: Heat-inactivated; TF: Transcription Factor; PFC: Prefrontal cortex; KEA3: Kinase Enrichment Analysis 3; CHEA3: ChIP-X Enrichment Analysis Version 3

### Active kinase and enriched transcription factor analysis

We then used KEA3 to identify active kinases responsible for the observed differential phosphorylation profiles. In male microglia, kinases such as NTRK2, PRKACA and PAK2 were predicted to be active, whereas in female microglia, kinases such as FGFR4, EPHA3 and MAP4K4 were predicted to be active; in male astrocytes, kinases such as MAPK1, TBK1 and PRKCD were predicted to be active, whereas in female astrocytes, kinases such as KSR1, PAK3 and NTRK1 were predicted to be active (Table 1). Protein kinases expressed as mRNA in mouse whole cortex and hippocampus (Figure 3) had minimal overlap with identified active kinases (Figure S1). These findings suggest kinase expression in male and female microglia or astrocytes poorly predicts kinase activity. Minimal overlap was also seen in active kinases (Table S2) between sexes (Figure 2C).

**Figure 3.**
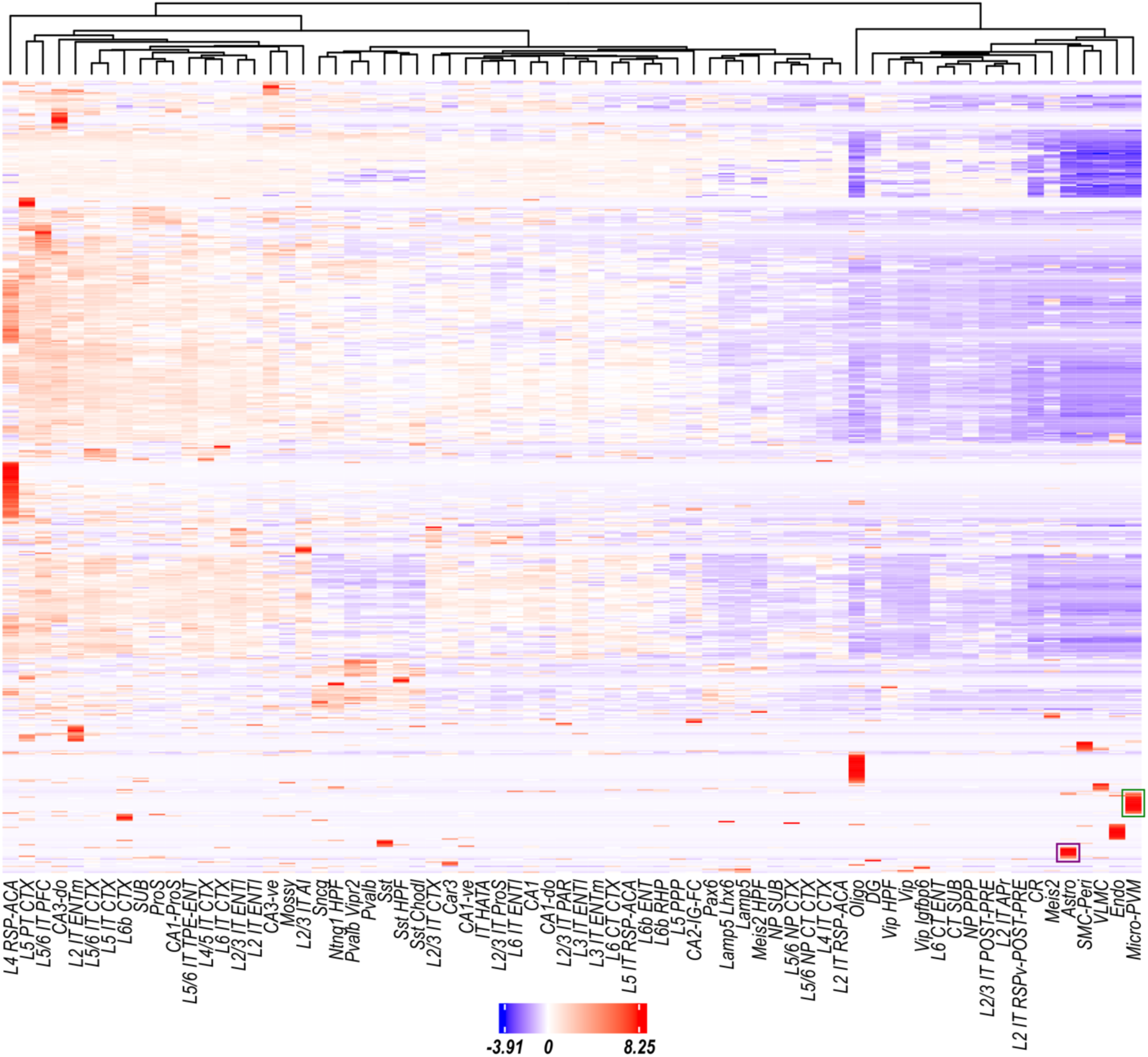
Cell-type specific kinase expression in mouse whole cortex and hippocampus. mRNA expression of the kinome in mice whole cortex and hippocampus was obtained from the Allen Brain Map 10x single-cell transcriptome reference atlas. Heatmap shows normalized mean gene expression of all kinases per cell type.

**Table 1.** Active kinases identified in female and male microglia or astrocytes. KEA3 was used to predict upstream active kinases responsible for the differential phosphorylation of reporter phosphopeptides observed in microglia or astrocytes from male and female wildtype mice PFC assayed on the PamGene PamStation®12 PamChip®4. Table shows uniquely identified kinases not identified in other groups. KEA3: Kinase Enrichment Analysis 3; PFC: Prefrontal cortex

| Kinase | M Micro | M Astro | F Micro | F Astro |
| --- | --- | --- | --- | --- |
| NTRK2 (neurotrophic receptor tyrosine kinase 2) | ✓ |  |  |  |
| PRKCA (protein kinase C alpha) | ✓ |  |  |  |
| PAK2 (p21 (RAC1) activated kinase 2) | ✓ |  |  |  |
| FYN (FYN proto-oncogene, Src family tyrosine kinase) | ✓ |  |  |  |
| PXK (PX domain containing serine/threonine kinase like) | ✓ |  |  |  |
| EPHB3 (EPH receptor B3) | ✓ |  |  |  |
| FGFR2 (fibroblast growth factor receptor 2) | ✓ |  |  |  |
| MAPK1 (mitogen-activated protein kinase 1) |  | ✓ |  |  |
| TBK1 (TANK binding kinase 1) |  | ✓ |  |  |
| PRKCD (protein kinase C delta) |  | ✓ |  |  |
| CAMKK1 (calcium/calmodulin dependent protein kinase kinase 1) |  | ✓ |  |  |
| NEK9 (NIMA related kinase 9) |  | ✓ |  |  |
| MAP3K20 (mitogen-activated protein kinase kinase kinase 20) |  | ✓ |  |  |
| MAPK14 (mitogen-activated protein kinase 14) |  | ✓ |  |  |
| RPS6KA3 (ribosomal protein S6 kinase A3) |  | ✓ |  |  |
| TYK2 (tyrosine kinase 2) |  | ✓ |  |  |
| STK4 (serine/threonine kinase 4) |  | ✓ |  |  |
| TRPM7 (transient receptor potential cation channel subfamily M member 7) |  | ✓ |  |  |
| FGFR4 (fibroblast growth factor receptor 4) |  |  | ✓ |  |
| EPHA3 (EPH receptor A3) |  |  | ✓ |  |
| MAP4K4 (mitogen-activated protein kinase kinase kinase kinase 4) |  |  | ✓ |  |
| LCK (LCK proto-oncogene, Src family tyrosine kinase) |  |  | ✓ |  |
| MAP3K4 (mitogen-activated protein kinase kinase kinase 4) |  |  | ✓ |  |
| PRKAA1 (protein kinase AMP-activated catalytic subunit alpha 1) |  |  | ✓ |  |
| STK38 (serine/threonine kinase 38) |  |  | ✓ |  |
| GSK3B (glycogen synthase kinase 3 beta) |  |  | ✓ |  |
| KSR1 (kinase suppressor of ras 1) |  |  |  | ✓ |
| PAK3 (p21 (RAC1) activated kinase 3) |  |  |  | ✓ |
| NTRK1 (neurotrophic receptor tyrosine kinase 1) |  |  |  | ✓ |
| AKT3 (AKT serine/threonine kinase 3) |  |  |  | ✓ |
| GRK1 (G protein-coupled receptor kinase 1) |  |  |  | ✓ |
| PDGFRA (platelet derived growth factor receptor alpha) |  |  |  | ✓ |
| PAK6 (p21 (RAC1) activated kinase 6) |  |  |  | ✓ |
| PAK4 (p21 (RAC1) activated kinase 4) |  |  |  | ✓ |
| ALK (ALK receptor tyrosine kinase) |  |  |  | ✓ |

We then used ChEA3 to identify transcription factors that are likely upstream of the active kinases and differentially phosphorylated peptides in each group. In male microglia, TFs such as NFATC1, E2F4 and SRF were predicted to be enriched, whereas in female microglia, TFS such as NR4A2, KLF7 and CSRNP2 were predicted to be enriched; in male astrocytes, TFs such as FOS, MEF2A and RFX5 were predicted to be enriched, whereas in female astrocytes, TFs such as ATF2, RORB and POU3F1 were predicted to be enriched (Table S2). Taken together, these TFs (Figure 2D), active kinases and differentially phosphorylated peptides indicated the presence of sex-specific signaling nodes in these cell types.

### Active signaling pathway analysis

We finally used the Kinograte package and the PCSF algorithm to reconstruct active signaling pathways by integrating differentially phosphorylated peptides, active kinases and enriched transcription factors in PPI networks. Pathway analysis of PPI nodes in male and female astrocytes or microglia revealed a total 1066 significantly enriched (FDR adjusted p-value < 0.05) Gene Ontology terms (Figure 4A). Functional clustering of enriched pathways with PAVER identified a total of 14 distinct pathway clusters in UMAP space (Figure 4D) showing varied enrichment across cell type and sex (Figure 4C) such as *regulation of cell differentiation*, *DNA-binding transcription factor binding*, and *neuron projection development* (Figure 4B). In male microglia, uniquely enriched active signaling pathways included *granulocyte differentiation*, *neurotrophin binding* and *regulation of DNA methylation*, whereas in female microglia, uniquely enriched active signaling pathways *included somatic recombination of immunoglobulin genes involved in immune response*, *T-helper 1 cell differentiation* and *negative regulation of glucocorticoid receptor signaling pathway*; in male astrocytes, uniquely enriched active signaling pathways included *p38MAPK cascade*, *regulation of extracellular exosome assembly* and *inflammatory response to wounding*, whereas in female astrocytes, uniquely enriched active signaling pathways included *neuronal ion channel clustering*, *nBAF complex* and *platelet-derived growth factor receptor binding* (Table 2). These findings indicated that these sex-specific active signaling pathways may lead to divergent functional outcomes in astrocytes and microglia.

**Figure 4.**
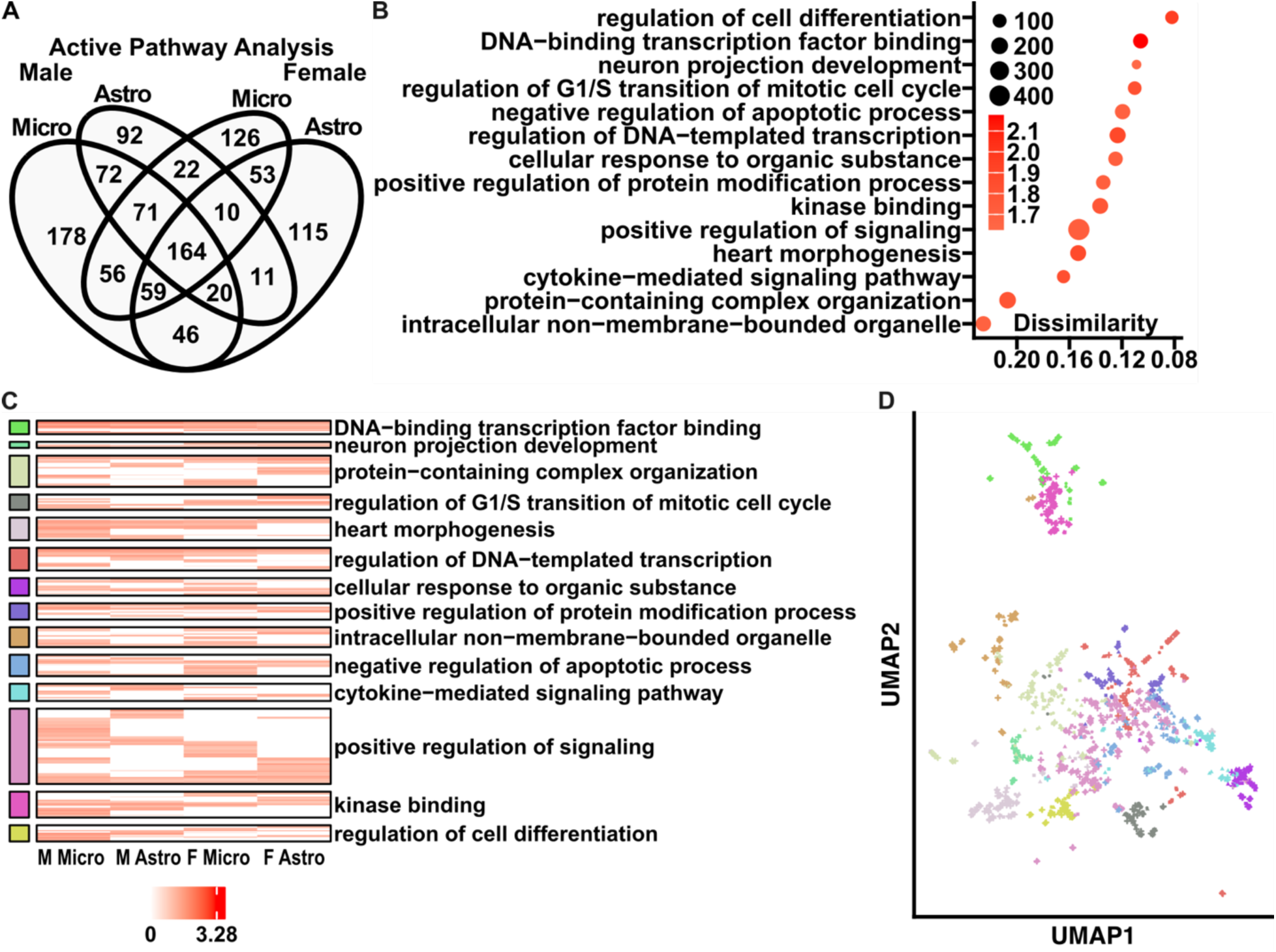
Functional interpretation of active signaling pathways enriched in female and male microglia or astrocytes. Nodes from PPI networks reconstructed by integrating differentially phosphorylated peptides, active kinases and enriched transcription factors identified in each group with PCSF were submitted to Enrichr for overrepresentation analysis and then clustered with PAVER, a meta-clustering method for pathways. **A.** Venn diagram overlap of significantly enriched (FDR adjusted p-value < 0.05) pathways identified in each group. **B.** Dot plot of functional pathway clusters identified by PAVER. Legend shows cluster mean log combined score and cluster size. X-axis shows dissimilarity of cluster MRT to its respective cluster. **C.** Heatmap of individual pathway enrichment and their respective cluster. Legend shows combined score. **D.** UMAP scatter plot of individual pathways colored by their respective cluster. PPI: protein-protein interaction; PCSF: Prize-Collecting Steiner Forest; PAVER: Pathway Analysis Visualization with Embedding Representations; MRT: Most Representative Term; UMAP: Uniform Manifold Approximation Projection

**Table 2.**
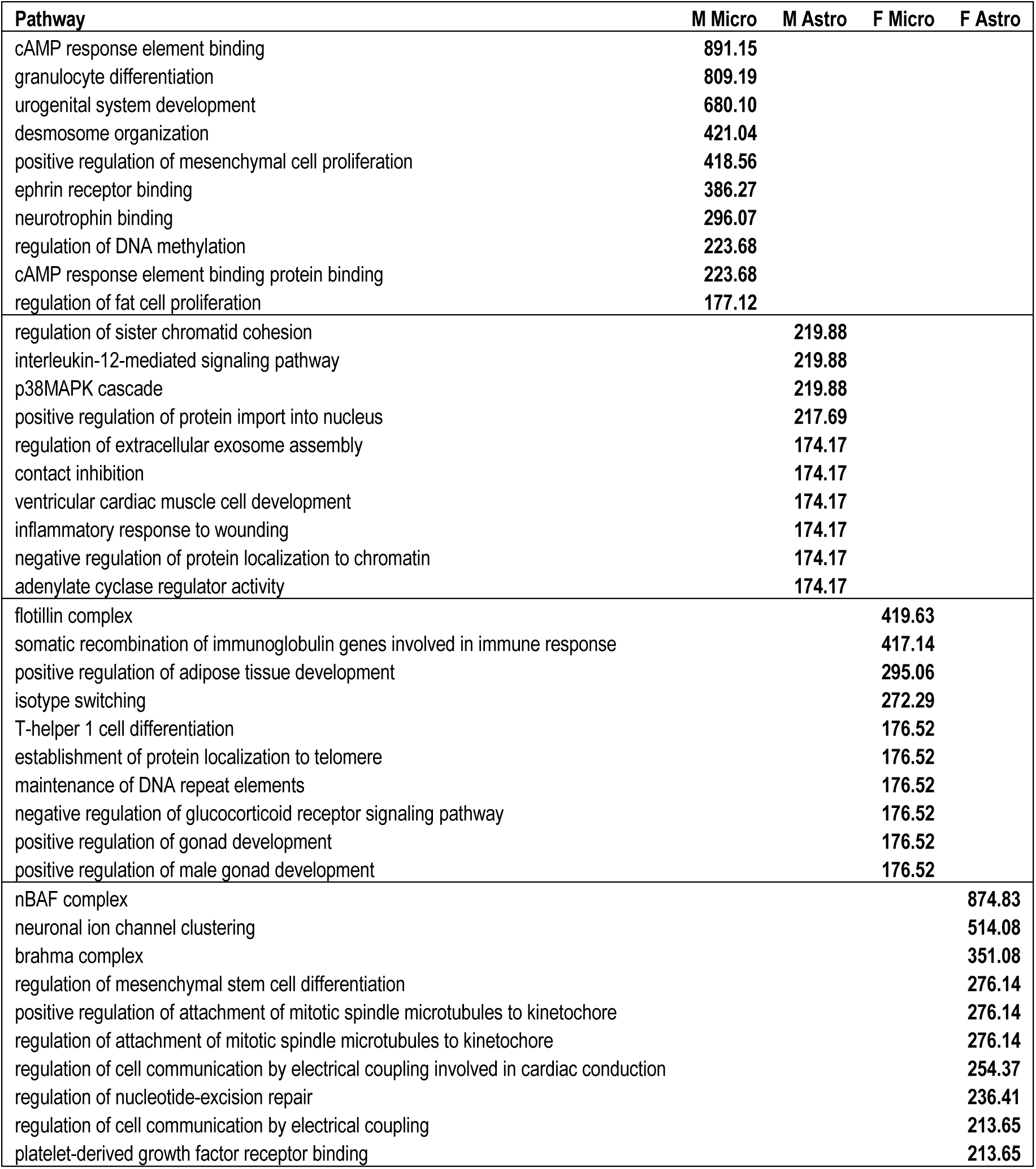
Active signaling pathways enriched in female and male microglia or astrocytes. Table shows combined scores of top ten unique significantly enriched (FDR adjusted p-value < 0.05) active signaling pathways identified in female and male microglia or astrocytes.

### Comparative analysis of cell and sex specific signaling networks

We visualized HGNC symbols annotated to active signaling pathways uniquely enriched in each group (Table S4). Comparative analysis revealed specific active signaling PPI networks present in both male and female microglia (Figure 5), and male and female astrocytes (Figure 6). In male microglia, the top nodes identified included PRKACA, ATF4 and SLC9A3R1, while in female microglia, the top nodes identified included TP53, ESR1 and RBBP6; in male astrocytes, the top nodes identified included TP53, FOS and ANXA2, while in female astrocytes, the top nodes identified included PRKACA, SLC9A3R1 and ITPR3 (Table 3). These findings indicated sex impacts active signaling nodes and networks in these cell types, and sex is an important biological variable in studies of brain cell signaling.

**Figure 5.**
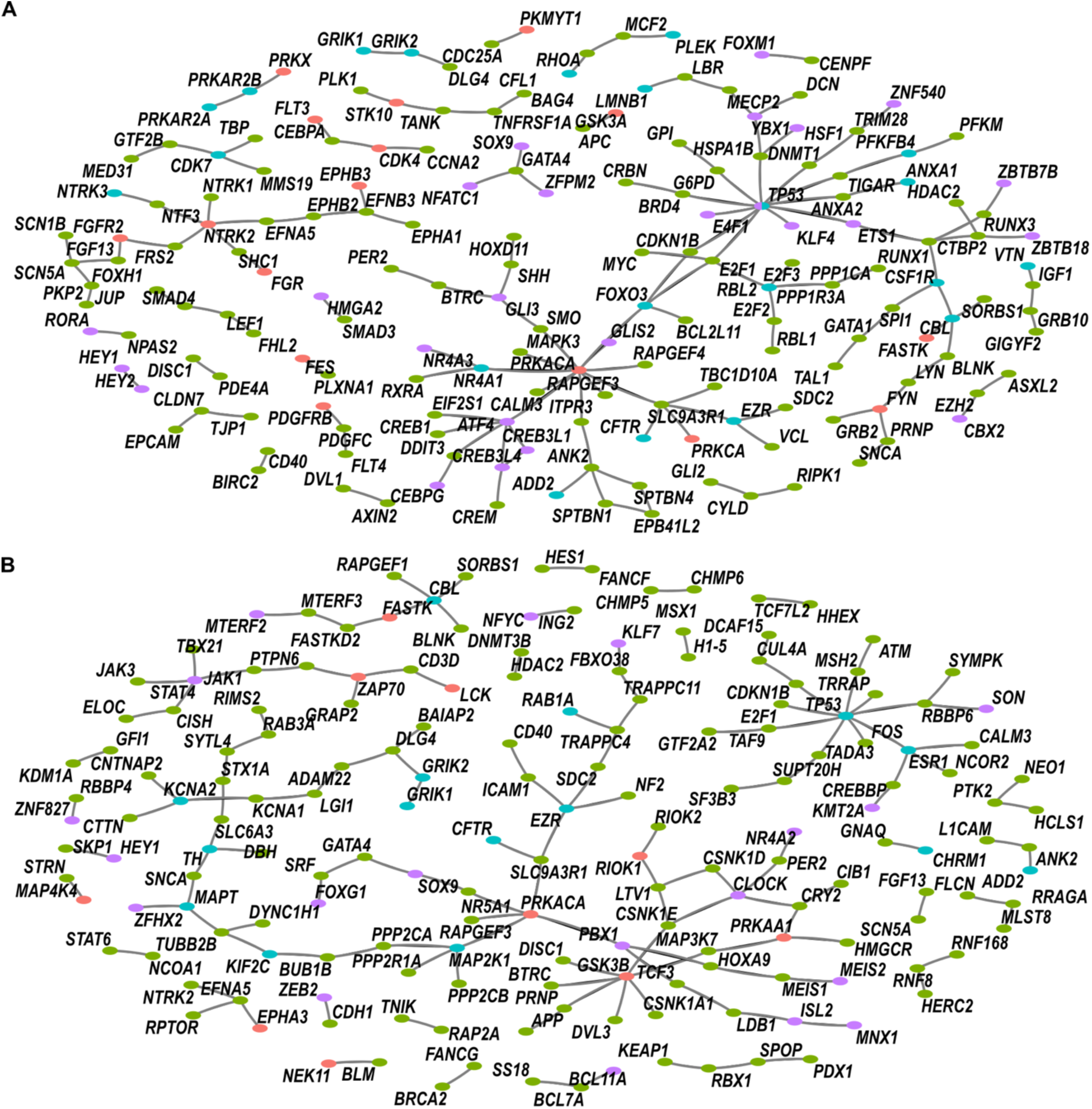
Sex-specific active signaling pathway networks enriched in male or female microglia. HGNC symbols annotated to active signaling pathways enriched uniquely in male or female microglia were used to generate PPI networks. Node color shows “hit” type (red: active kinase, purple: upstream transcription factor, blue: differentially phosphorylated peptide, green: Steiner hidden node). **A.** Male microglia active signaling network. **B.** Female microglia active signaling network. HGNC: HUGO Gene Nomenclature Committee; PPI: protein-protein interaction

**Figure 6.**
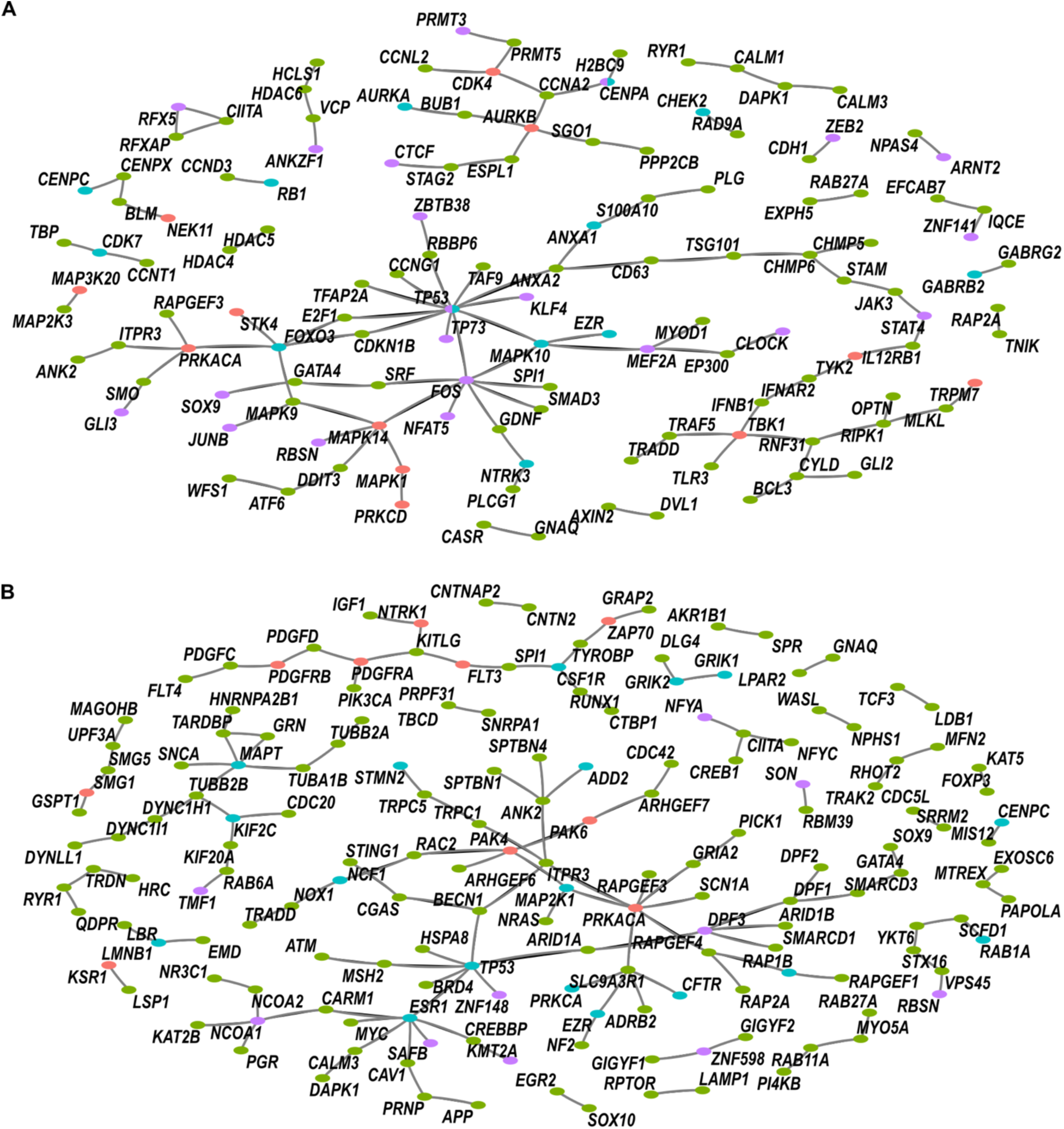
Sex-specific active signaling pathway networks enriched in male or female astrocytes. HGNC symbols annotated to active signaling pathways enriched uniquely in male or female astrocytes were used to generate PPI networks. Node color shows “hit” type (red: active kinase, purple: upstream transcription factor, blue: differentially phosphorylated peptide, green: Steiner hidden node). **A.** Male astrocytes active signaling network. **B.** Female astrocytes active signaling network. HGNC: HUGO Gene Nomenclature Committee; PPI: protein-protein interaction

**Table 3.** Interacting nodes in active signaling networks identified as enriched in male and female astrocytes or microglia. Table shows the top ten nodes by hub score (eigencentrality influence) in active signaling PPI networks identified as enriched in male and female astrocytes or microglia. PPI: protein-protein interaction

| Node | M Micro | M Astro | F Micro | F Astro |
| --- | --- | --- | --- | --- |
| PRKACA (protein kinase cAMP-activated catalytic subunit alpha) | 1 |  |  | 1 |
| ATF4 (activating transcription factor 4) | 0.449 |  |  |  |
| SLC9A3R1 (Sodium-hydrogen antiporter 3 regulator 1) | 0.401 |  |  | 0.647 |
| FOXO3 (forkhead box O3) | 0.372 |  |  |  |
| NR4A1 (nuclear receptor subfamily 4 group A member 1) | 0.349 |  |  |  |
| ITPR3 (inositol 1,4,5-trisphosphate receptor type 3) | 0.322 |  |  | 0.481 |
| SMO (smoothened, frizzled class receptor) | 0.32 |  |  |  |
| CALM3 (calmodulin 3) | 0.29 |  | 0.231 |  |
| GLIS2 (GLIS family zinc finger 2) | 0.29 |  |  |  |
| RAPGEF4 (Rap guanine nucleotide exchange factor 4) | 0.29 |  |  | 0.456 |
| TP53 (tumor protein p53) |  | 1 | 1 |  |
| FOS (Fos proto-oncogene, AP-1 transcription factor subunit) |  | 0.703 | 0.366 |  |
| ANXA2 (annexin A2) |  | 0.35 |  |  |
| RBBP6 (RB binding protein 6, ubiquitin ligase) |  | 0.345 | 0.425 |  |
| KLF4 (KLF transcription factor 4) |  | 0.312 |  |  |
| TAF9 (TATA-box binding protein associated factor 9) |  | 0.312 | 0.422 |  |
| CCNG1 (cyclin G1) |  | 0.312 |  |  |
| TFAP2A (transcription factor AP-2 alpha) |  | 0.312 |  |  |
| TP73 (tumor protein p73) |  | 0.312 |  |  |
| MAPK14 (mitogen-activated protein kinase 14) |  | 0.303 |  |  |
| ESR1 (estrogen receptor 1) |  |  | 0.633 |  |
| E2F1 (E2F transcription factor 1) |  |  | 0.366 |  |
| TRRAP (transformation/transcription domain associated protein) |  |  | 0.366 |  |
| CREBBP (CREB binding protein) |  |  | 0.267 |  |
| NCOR2 (nuclear receptor corepressor 2) |  |  | 0.231 |  |
| MAP2K1 (mitogen-activated protein kinase kinase 1) |  |  |  | 0.475 |
| RAPGEF3 (Rap guanine nucleotide exchange factor 3) |  |  |  | 0.342 |
| SCN1A (sodium voltage-gated channel alpha subunit 1) |  |  |  | 0.342 |
| GRIA2 (glutamate ionotropic receptor AMPA type subunit 2) |  |  |  | 0.254 |
| PAK4 (p21 (RAC1) activated kinase 4) |  |  |  | 0.226 |
| EZR (ezrin) |  |  |  | 0.226 |

## Discussion

In this study, we sought to delineate sex-specific differences in active signaling pathways in astrocytes and microglia isolated from the prefrontal cortex (PFC) of male and female mice. Using the PamGene PamStation®12 kinome activity profiling platform, we measured kinase activity across complex signaling networks, providing an unprecedented atlas of active pathways in these brain cells. We ultimately identified critical cell and sex-specific alterations in these pathways that could have significant implications for understanding variability in brain diseases and their treatments.

Our analysis demonstrated substantial differences in kinase activity between male and female astrocytes and microglia, with distinct phosphorylation patterns of reporter phosphopeptides observed in each group. Among the kinases identified as active, several were unique to specific cell types and sexes. In male microglia, kinases such as NTRK2, PRKACA and PAK2 were prominently active, indicative of their roles in mediating neuroinflammation and cellular proliferation [49–51]. In contrast, female microglia were characterized by the activity of kinases such as FGFR4, EPHA3 and MAP4K4, which are involved in immune response regulation and cellular differentiation [52–54]. In astrocytes, male-specific active kinases included MAPK1, TBK1 and PRKCD, which are associated with synaptic plasticity and inflammatory responses [55–57]. Female astrocytes conversely showed activity in kinases such as KSR1, PAK3 and NTRK1, which are implicated in neurogenesis and synaptic modulation [58–62]. The differential activation of these kinases suggests that sex-specific factors influence how microglia or astrocytes contribute to brain function, which could impact the onset and progression of brain disease. Our analysis also showed distinct sex-specific transcription factor enrichment patterns in both astrocytes and microglia. In male microglia, transcription factors such as NFATC1, E2F4 and SRF were enriched, these having roles in immune regulation and cell cycle control [63–65]. Female microglia, however, showed enrichment for transcription factors like NR4A2, KLF7 and CSRNP2, which are involved in cellular differentiation and stress responses [66–68]. In astrocytes, male-specific transcription factors included FOS, MEF2A and RFX5, which are known to play roles in synaptic regulation and neuronal development [69–71]. Female astrocytes, enriched for transcription factors such as ATF2, RORB and POU3F1, are involved in processes related to neuroprotection and circadian rhythm regulation [72]. These findings further support the notion that male and female microglia operate under functionally distinct active signaling networks.

The integration of kinase activity data and transcription factor analysis with protein-protein interaction networks provided a comprehensive view of the active signaling pathways in male and female astrocytes and microglia. Functional clustering identified 14 distinct pathway clusters, including those related to *cell differentiation*, *DNA-binding transcription factor activity* and *neuron projection development*. These clusters showed varied enrichment across cell types and sexes, suggesting that these sex-specific signaling pathways could be driving different functional outcomes seen in astrocytes and microglia [73–75].

We also identified unique active signaling pathways in each group, revealing that sex influences signaling network architecture. For example, male microglia showed enrichment in pathways related to granulocyte differentiation and neurotrophin binding, which are crucial for immune responses and neuronal survival [76, 77]. Female microglia were enriched in pathways related to immune modulation and glucocorticoid signaling, which may impact their response to stress and injury [78, 79]. In astrocytes, male-specific pathways included those involved in the p38MAPK cascade and inflammatory responses, which are critical for regulating neuroinflammation [80], while female-specific pathways were related to neuroprotection and synaptic organization, which may impact their response to disease processes by enhancing cellular resilience and supporting synaptic integrity [78, 79]. The identification of these pathways provided sex-specific targets that could modulate microglia and astrocyte function. Our findings also have significant implications for understanding the molecular basis of sex differences in brain diseases. The distinct kinase activity patterns observed in male and female astrocytes and microglia could contribute to sex-specific susceptibility and progression of neurodegenerative disorders such as Alzheimer’s disease and other neuroinflammatory conditions [81]. For instance, the differential activation of kinases like PRKACA and MAPK1 in male and female astrocytes, as well as the activation of kinases such as FGFR4 and EPHA3 in female microglia and NTRK2 and PAK2 in male microglia, may influence the development of neurodegenerative pathology [82, 83]. This suggests considering sex as a biological variable in therapeutic development is highly important, as sex-specific active kinase signaling could lead to different disease mechanisms and treatment responses in males and females. Interestingly, we found there was minimal overlap between protein kinases expressed as mRNA versus those predicted to be active in each cell type. This aligns with the generally poor correlation seen between gene expression and protein abundance, or between protein abundance and protein activity [16, 18]. Taken together, in light of transcription being thought of as an “undruggable” target [84], and over 3,000 unsuccessful clinical trials targeting gene-level changes for neurodegenerative disease [85], the sum of our findings suggests studying active kinase signaling profiles in brain disease as a promising research space for drug discovery. Despite the valuable insights gained from this study, several limitations should be acknowledged. The PamGene PamStation®12 platform, while well-established for measuring kinase activity, relies on peptide-based assays that may not capture the full complexity of in vivo kinase-substrate interactions. Future studies with different animal models or human samples, larger sample sizes and additional validation techniques, such as mass spectrometry-based phosphoproteomics, will be needed to confirm and extend our results. Such studies could help confirm the sex-specific differences in kinase activity and identify species-specific or disease-related variations.

In conclusion, our study provides a comprehensive analysis of sex-specific differences in active kinase signaling pathways in astrocytes and microglia. These findings emphasize the importance of considering sex and functional network activity as a critical factor in brain research and therapy development. By advancing our understanding of the molecular mechanisms underlying brain disease, this study contributes to the growing field of sex-specific medicine and offers a framework for improving brain health.

## Acknowledgements

This work was supported by NIH NIGMS T32-G-RISE grant number 1T32GM144873-01, NIH NIMH grant number R01MH107487, NIH NIMH grant number R01MH121102, and NIH NIA grant number R01AG057598, and NIH NIA grant number R01AG083628.

