## Supplemental Materials for "Characterization of active kinase signaling pathways in astrocytes and microglia"

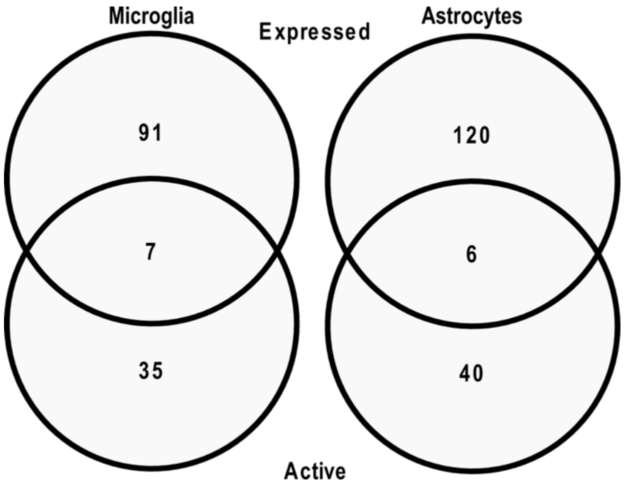


**Figure S1. Kinases expressed and/or active in microglia or astrocytes.**

Venn diagram shows overlap of kinases expressed as mRNA versus kinases determined as active in microglia or astrocytes.

**Table S1. Differential phosphorylation of reporter phosphopeptides.**

Peptide table obtained from the KRSA software shows log fold changes of peptides measured on the PamGene PamStation®12 PamChip®4 in each group versus their respective control.

KRSA: Kinome Random Sampling Analyzer

**Table S2. Kinase enrichment analysis using KEA3.**

Table shows complete kinase enrichment results using differentially phosphorylated peptides as input to KEA3's kinase-substrate interaction libraries.

KEA3: Kinase Enrichment Analysis 3.

**Table S3. Transcription factor enrichment analysis using CHEA3.**

Table shows transcription factor enrichment results using differentially phosphorylated peptides and active kinases as input to CHEA3's brain-specific transcription factor library.

CHEA3: ChIP-X Enrichment Analysis Version 3

**Table S4. Functional clustering of active signaling enriched pathways in male and female astrocytes or microglia.**

Table shows log combined scores of pathways in each group and their associated cluster identified by PAVER.

PAVER: Pathway Analysis Visualization with Embedding Representations
